# Population and adaptation history of 739 *Thlaspi arvense* natural accessions

**DOI:** 10.1101/2025.03.21.644658

**Authors:** Xing Wu, Ruth Epstein, Maliheh Esfahanian, Barsanti Gautam, Marcus Griffiths, Jules Perez, Kerrie Barry, Anna Lipzen, Chris Daum, Yuko Yoshinaga, Chanaka Roshan Abeyratne, Zhaslan Akhmetov, Sebastian Toro Arana, Ryan Bayliss, Bhabesh Borphukan, Anthony Brusa, Hari Chhetri, Rachel Combs-Giroir, Lucas Czech, Marcin K. Dyderski, Eva Serena Gjesvold, Grzegorz Grzejszczak, Shannon Hateley, Nicholas Heller, Danielle Hoffmann, Nikhil Jaikumar, Brice A. Jarvis, Vanessica Jawahir, Marcin Klisz, Peter Kruse, Matthew Lane, Arjuman Lima, Alexander Liu, Gabriela Madrid, Maggie Marlino, Michaela McGinn, Mirko Pavicic, William Perry, Manesh Shah, Jason Thomas, Alice Townsend, Thiranya L. Wanigarathna, Tad Wesley, Bryan Connolly, Yong Pyo Lim, Radosław Puchałka, Alexander Wirth, Andrea R. Gschwend, Pubudu P. Handakumbura, Daniel Jacobson, Dmitri A. Nusinow, Seung Y. Rhee, Karen A. Sanguinet, Christopher N. Topp, Jeremy Schmutz, M. David Marks, Winthrop Phippen, Ratan Chopra, John C. Sedbrook, Moisés Expósito-Alonso

## Abstract

Pennycress (*Thlaspi arvense*) is a promising intermediate oilseed crop, producing oil suitable for conversion to biofuels—including aviation fuels. While domestication efforts are ongoing, a deeper understanding of the genetic architecture of traits is crucial for informing future breeding efforts. Here, we conducted the largest genomic and phenotypic survey of pennycress to date, analyzing 739 accessions collected across four continents. Leveraging whole-genome sequencing and field-collected phenotypes, we characterized the standing genetic variation underlying key agronomic traits and climate resilience. Our findings revealed multiple independent migration events to North America, with substantial genetic admixture. We identified homologs of *Arabidopsis thaliana* flowering-time genes that contribute to adaptation and demonstrated the agronomic benefits of winter-type pennycress. Furthermore, through multi-season field trials, we identified a genomic region containing a cluster of *mTERF* genes strongly associated with green canopy coverage, a critical trait for biomass retention and yield stability. These insights provide a genomic roadmap for accelerating pennycress domestication and improving its resilience to climate variability.

## Introduction

Field pennycress, *Thlapsi arvense L.* (Brassicaceae), is a ruderal species being targeted for domestication for use as an intermediate oilseed crop. In the lower Midwestern United States, pennycress has a short enough life cycle to fit in the fallow season within corn and soybean rotations, resulting in three crop yields in two years ^1–4^. Similar to the wild progenitors of major crops ^5^, pennycress naturally possesses desirable traits, such winter hardiness, high grain yield, a short seed to seed cycle, a potential to diversify ecosystems, and annuality, which favors it as a candidate for neo-domestication as a new crop ^6–8^. Pennycress seeds also contain up to 35% oil in triacylglycerol, which is well suited for conversion to renewable diesel and sustainable aviation fuel (SAF) ^9–11^. In addition to high oil content, pennycress seeds contain over 20% edible protein content, which makes the meal suitable for use as animal feed^12^.

Despite its innate useful traits, pennycress must undergo major genetic and morphological changes for domestication such as the adoption of earlier flowering, non-shattering seed pods, increased seed oil content, reduced glucosinolate content, reduced weediness, and enhanced tolerance to abiotic stresses^1,11,13–17^. Work to domesticate pennycress and deploy it as a new oilseed crop is well underway^1,11,18,19^. Forward genetic screens have successfully utilized near-infrared spectroscopy to identify mutant lines with desirable seed composition, offering valuable insights for future breeding efforts^20^. Reverse genetics approaches have also been effectively employed to modify seed composition for renewable fuel production, reduce weediness, improve meal quality, and improve environmental sensing traits, contributing to the creation of a new crop^21,22^.

Historically, the domestication of major crops involved thousands of years of selective breeding, however, neo-domestication efforts can be accelerated with conventional methods such as chemical mutagenesis along with new technologies like marker-assisted breeding and gene editing^1,10,17,19,23^. Even here, pennycress has another advantage: It is genetically closely related to the model plant, *Arabidopsis thaliana (A. thaliana)* ^18,24^. Like *A. thaliana*, pennycress can be transformed easily via a floral dip method, is primarily self-fertilized, and has a relatively small diploid genome^18,19,25^. In addition, pennycress has an almost 1-to-1 orthologous gene correspondence with *A. thaliana,* suggesting a conserved genetic architecture for most heritable traits^18^. Importantly, unlike many other Brassicaceae species that have undergone extra rounds of whole genome duplication introducing functional redundancy^26^, the plethora of scientific knowledge of *A. thaliana* can be translated into creating desirable knockout or knockdown mutations in pennycress to advance domestication efforts.

To inform future breeding efforts, the full spectrum of natural genetic variation in pennycress must be explored and understood^23,27^. Here, we present the first large-scale genomic analysis of 739 diverse pennycress accessions, uncovering its complex evolutionary history and the genetic basis for growth type and climate adaptation. We identified genetically distinct subpopulations in Asia and evidence for multiple migration events into North America with substantial admixing amongst European and North American samples. Genome-wide association studies (GWAS) linked growth type to key, large effect *A. thaliana* homologous flowering regulators, *DELAY OF GERMINATION 1* (*DOG1*) and *FLOWERING LOCUS C* (*FLC*), while climate adaptation exhibited a more polygenic architecture. A large-scale field experiment further identified a significant association near mitochondrial transcription termination factor 5 (*MTERF5*) with green canopy coverage, highlighting a potential gene of interest for crop improvement. Together, our findings provide a foundational genomic resource and novel insights into pennycress evolution, informing its future improvement as a climate-resilient cover crop.

## Results

### Population structure reveals multiple migrations and admixing events

To characterize the extent and distribution of standing genetic diversity in our pennycress population, we analyzed whole-genome sequencing (WGS) data from 739 accessions, including 479 newly sequenced in this study and 260 from previously published datasets (**Table S1**)^28–30^. The new accessions were collected by the Integrated Pennycress Resilience Project (IPReP) Consortium and citizen scientists from diverse ecological settings across 20 countries spanning four continents—Europe, Asia, North America, and South America—with precise geographic coordinates recorded for each sample (**Figure 1A**). WGS was performed at an average depth of 24.1X, yielding 4.6 TB of total raw data. After filtering out low-quality sequencing reads, we aligned reads to the pennycress reference genome^29^, achieving an average mapping rate of 94.1%. Using an established plant genomic variant discovery pipeline^31^, we identified 5,428,817 high-confidence single nucleotide polymorphisms (SNPs), ∼1% of the pennycress genome, including 211,466 located in coding regions, resulting in 107,219 nonsynonymous mutations and 3,965 early stop codons. In addition, we detected 3,375,371 high-confidence indels, 64,697 of which were positioned within coding sequences (**Table S2**).

**Figure 1.**
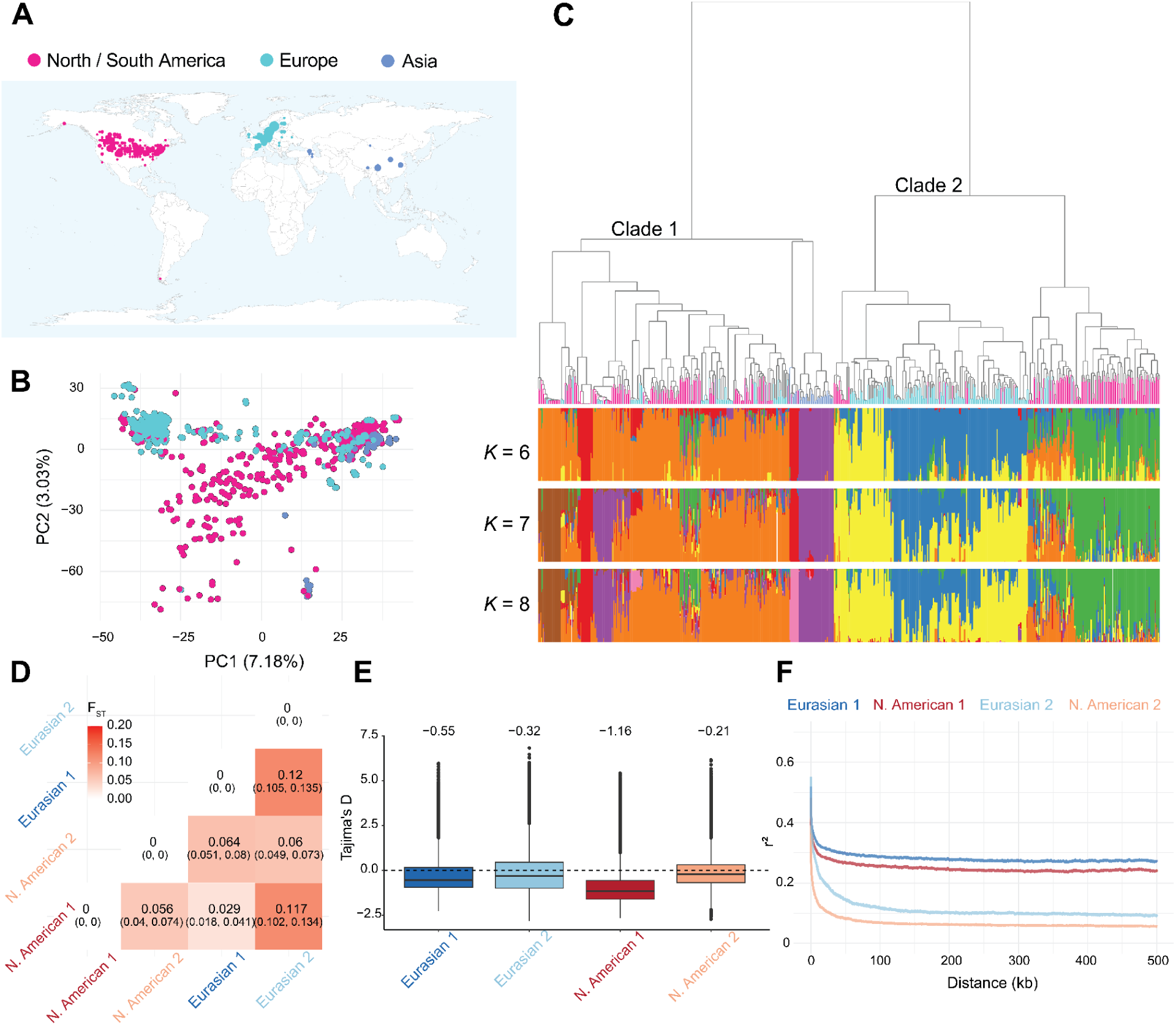
Genetic diversity and population structure of a diverse pennycress collection. **(A)** A world map showing the collection sites of 739 natural pennycress accessions. **(B)** Principal component analysis (PCA) of 739 accessions using LD-pruned genetic variants. **(C)** Hierarchical clustering of 739 accessions based on identity-by-state (IBS) genetic distance (top), alongside population structure analysis (bottom). **(D)** Pairwise F_ST_ between the 4 sub-clades with 95% confidence intervals. Clade 1 includes N. American 1 and Eurasian 1, while Clade 2 includes N. American 2 and Eurasian 2. **(E)** Genome-wide Tajima’s D values within each sub-clade as defined by IBS genetic distance. **(F)** Genome-wide LD decay across different sub-clades. Colored lines correspond to LD decay within each sub-clade.

Armed with the largest sequenced pennycress population to date, we first aimed to investigate patterns of genetic diversity across space. We performed a principal component analysis (PCA) using 5.4 million high-confidence SNPs, revealing significant divergence among accessions from different continents with the first two principal components accounting for 7.18% and 3.03% of the total genetic variance, respectively (**Figure 1B**). PC1 primarily separated European populations while PC2 distinguished Asian populations. Most notably, several North American samples closely cluster with either Asian or European collected accessions, while other North American samples seem to be genetically diverged from any Eurasian sample, leading to an investigation into the genetic relationships between our pennycress samples.

To further explore the genetic structure of our population revealed by the PCA, we conducted hierarchical clustering of our 739 accessions based on identity-by-state (IBS) distance. This analysis revealed two major clades, suggesting two distinct ancestries in the demographic history of pennycress (**Figure 1C**). Clade 1 encompassed all Asian accessions, and included interspersed North American and European accessions. On the other hand, Clade 2 consisted exclusively of North American and European accessions, which were distinctly separated within the clade (**Figure 1C**). Since there was a separation between North American and European samples in Clade 2, we decided to further explore their genetic divergence, and calculated the fixation index (F_ST_), both between and within the two clades (**Figure 1D**). North American accessions in Clade 2 were significantly differentiated from its Eurasian counterpart (F_ST_ = 0.06, 95% CI: 0.049-0.073). In contrast, North American Clade 1 samples and Eurasian Clade 1 samples had the lowest genetic differentiation (F_ST_ = 0.029, 95% CI: 0.018-0.041), suggesting a very recent migration to North America.

Given these varying levels of genetic differentiation, we sought to investigate whether these sub-clades have been evolving neutrally. To assess this, we performed a genome-wide calculation of Tajima’s D^32^. Our analysis revealed that the North American Clade 1 had a low Tajima’s D value of −1.16, suggesting a recently experienced selective sweep or a population bottleneck (**Figure 1E**). To explore the possibility of a bottleneck, we estimated the effective population size (N_e_) of sub-clades in the recent past, which can be used to identify past bottlenecks (reducing N_e_), population expansions (increasing N_e_), and divergence times (diverging N_e_). Using a site frequency spectrum (SFS)-based method to estimate N_e_ ^33^, we found that all clades have a small N_e_, likely due to high homozygosity caused by the self-fertilizing preference of pennycress. Nevertheless, we found the North American Clade 1 to have the lowest N_e_ in the recent past out of all sub-clades (**Figure S1**), suggesting it might be currently undergoing a bottleneck or is in population decline. Corroborating this, we found that North American Clade 1 samples had the lowest percentage of heterozygous sites, 0.75% (**Figure S2**).

After finding two distinct ancestries in pennycress, we performed model-based clustering to identify if there is a fine-scale population structure embedded within our larger population (**Figure 1C**). We not only observed past admixture events between European accessions (blue and yellow) in Clade 2 but also between North American accessions from Clades 1 and 2 (green and orange) (**Figure S3**). Within North America, there is a zone of admixing in the Midwestern United States, resulting in individuals with both green and orange, with the green sub-group occupying mostly Southern parts of the US, while the orange sub-groups occupy Northern parts of the US and Canada (**Figure S3**). Groups that show evidence of substantial admixture (**Figure 1C**) have the lowest amount of genetic differentiation, for instance, European samples in Clade 2 (blue and yellow) have a low F_ST_ (F_ST_ = 0.034, 95% CI: 0.02-0.053, **Figure 1D, Figure S4**). Nevertheless, despite evidence of substantial admixing, we still identified distinct subpopulations, such as the Asian population of Clade 1 which has two subpopulations: the Chinese (purple) and Armenian (red) groups (**Figure 1C**). The Armenian population is highly differentiated from the rest of the population (median F_ST_ = 0.191) and potentially had a recent population contraction given its high Tajima’s D of 2.76 (**Figure S4**).

Lastly, we assessed the rate of linkage disequilibrium (LD) decay because it can provide insight into genome-wide recombination rates and can determine the mapping resolution for GWAS in diverse accessions. Thus, we calculated LD decay across 739 accessions and examined LD decay in each of the four sub-clades and seven subpopulations (**Figure 1F, S4**). Overall, pennycress exhibits relatively large LD blocks, with r^2^=0.21 at a distance of 100 kb. Among our 4 sub-clades, North American and Eurasian Clade 1 samples display the slowest LD decay, maintaining r^2^>0.25 beyond 500 kb, while the North American and Eurasian Clade 2 samples exhibited the most rapid decay, with r^2^<0.1 after 100 kb. Interestingly, the North American samples, regardless of clade membership, experiences more rapid LD decay than its Eurasian counterparts supporting the idea that these samples might have different rates of recombination and/or divergent demographic histories.

### Genome-wide signals of natural selection in North America accessions

As the major goal of pennycress is to establish it as a cover crop grown primarily in the US Midwest, we wanted to examine if any of our North American-collected samples have adapted to a North American climate. If adaptation has occurred, we would expect to find evidence of selective sweeps—events where strong selection at a specific locus drives nearby alleles to fixation, resulting in a distinct pattern of homozygosity. To investigate selective sweeps both within and between the two clades, we applied the Cross-Population Composite Likelihood Ratio (XP-CLR) test, which models multilocus allele frequency differentiation between populations^34^. The XP-CLR test between the North American & Eurasian Clade 1 samples showed no detectable selective sweeps (**Figure 2A**). In comparison, North American and Eurasian Clade 2 had a few candidate regions for selective sweeps, with a significant peak on chromosome 2 spanning 78 kb (**Figure 2B**). Within this peak on chromosome 2, we identified 20 genes, including a DNA-directed RNA polymerase subunit, an asparagine synthase, two cyclin-dependent kinases, and most interestingly, a cluster of nine MADS-Affecting Flowering (MAF) transcription factors (**Figure 2C**), known to be involved in the regulation of flowering time in *A. thaliana* and in *Arabis* species ^35,36^. Although North American Clade 2 is more differentiated from its Eurasian counterpart, in comparison to Clade 1, we identify very few selective sweeps, suggesting adaptation to North America has not occurred or is just beginning.

**Figure 2.**
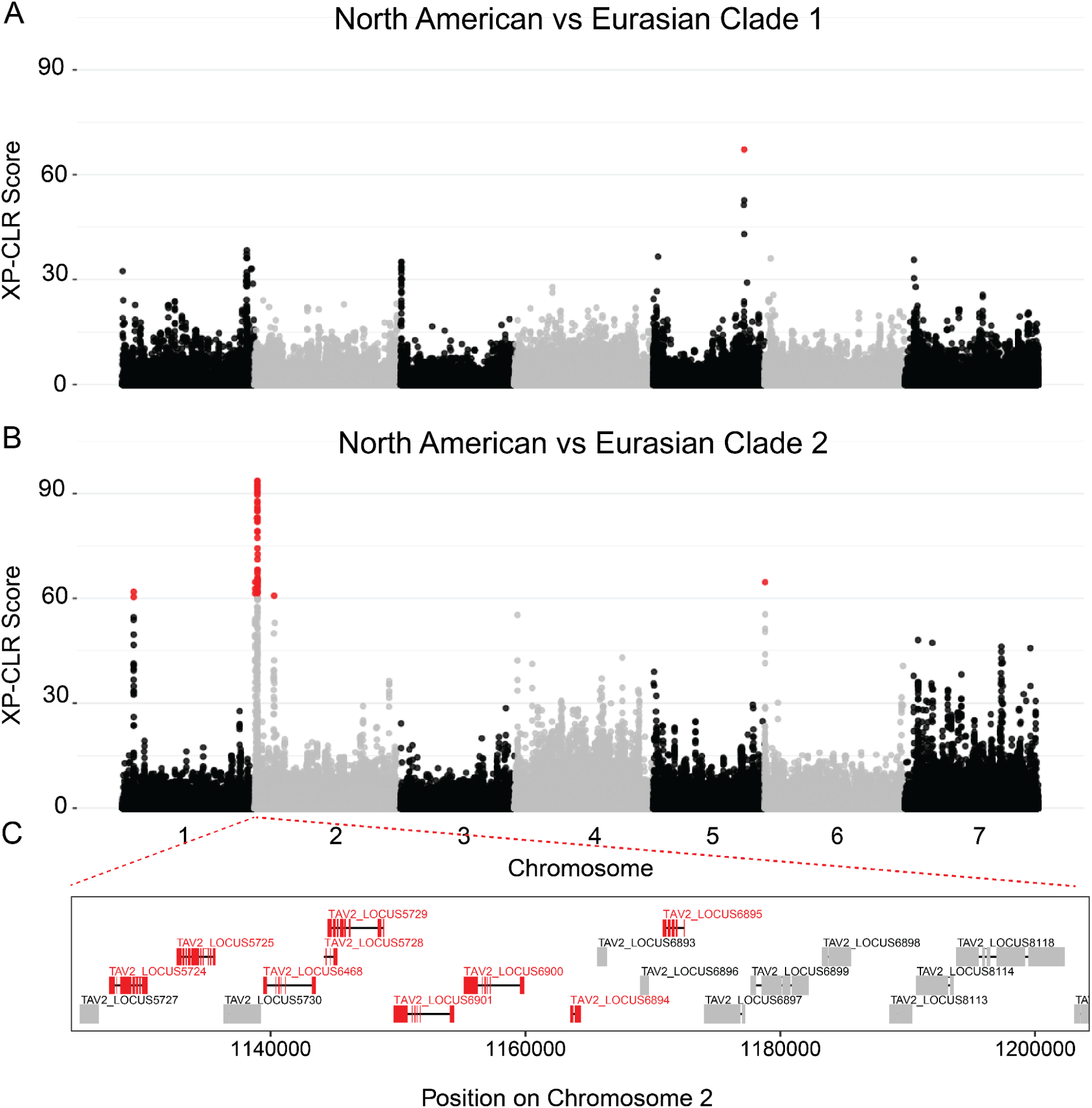
Manhattan plot of XP-CLR between: **(A)** North American Clade 1 & Eurasian Clade 1, **(B)** North American Clade 2 & Eurasian Clade 2. Red dots signify windows with a XP-CLR score above 60, which represents the 99th percentile of scores for both comparisons.

### Genetic architecture of growth type

Field pennycress exhibits two distinct growth types: spring and winter. Spring-type pennycress does not require cold temperatures to induce flowering (vernalization) and therefore can complete its life cycle within a single growing season. Winter-type pennycress requires vernalization before flowering which favors germination in the fall, resulting in a rosette overwinter before flowering in the following spring^8^. In this study, we phenotypically characterized the growth type of 453 natural accessions through multiple field seasons and detailed observations (**Materials & Methods**), categorizing them as spring, winter, or mixed growth types. Among these accessions, 172 were classified as spring type, 247 as winter type, and 34 as mixed type.

To elucidate the genetic basis of growth type in field pennycress, we conducted a genome-wide association study (GWAS) on 419 accessions, excluding the 34 mixed-type accessions. The narrow-sense heritability (h^2^) of growth type was estimated at 0.503 (+/− 0.067), highlighting a strong genetic contribution to this trait. Using GEMMA and HapFM, we identified significantly associated peaks on chromosomes 2 and 6, with both methods detecting associations in the same genomic regions (**Figure 3A, S5**). The chromosome 2 peak spanned 382 kb, 38 genes, and includes TAV2_LOCUS5510, homologous to AT5G45830, *DELAY OF GERMINATION (DOG1)*, in *A. thaliana*. The chromosome 6 peak spanned 763 kb with 168 genes, and includes TAV2_LOCUS21312, homologous to AT5G10140, *FLOWERING LOCUS C* (*FLC*), in *A. thaliana.* Additionally, HapFM identified a significant locus on chromosome 3 spanned 103 kb with 8 genes, encompassing a cluster of duplicated genes homologous to *SUPPRESSOR OF drm1 drm2 cmt3* (*SDC*) in *A. thaliana*.

**Figure 3.**
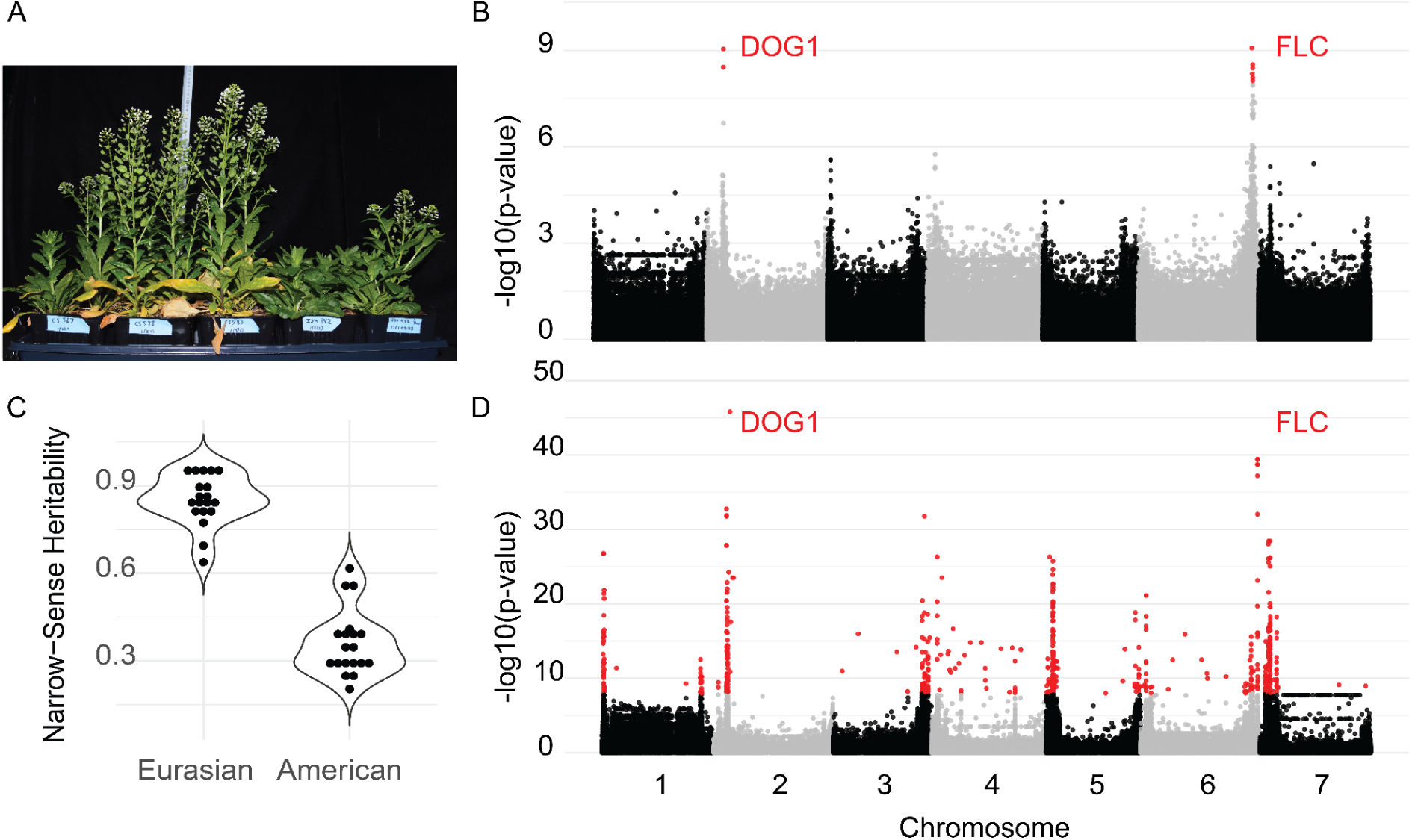
GWAS for growth type and BIOCLIM variables. (**A**) Growth chamber-grown plants which had been vernalized at least 21 days at 4 °C as seedlings to induce flowering of winter-types. Note the natural variation in flowering time. **(B)** The Manhattan plot of growth-type GWAS on 419 pennycress accessions. **(C)** The distribution of narrow-sense heritability of 19 BIOCLIM variables between Eurasian and North American populations **(D)** The Manhattan plot of the annual mean temperature of the Eurasian population.

### Genetic architecture of climate adaptation

Many plant populations exhibit local adaptation to their native climates—a phenomenon known as climate adaptation—often driven by standing genetic variation within the species^37–39^. In the PCA analysis of the 739 pennycress accessions, we found PC1 to be significantly associated with the minimum temperature of the coldest month at each collection site (Pearson’s correlation, *r* = −0.393, *P* = 2.2 × 10^−16^ **Figure S6**), suggesting climate adaptation plays a role in shaping genetic differentiation across the natural range of pennycress.

To investigate the genetic basis of climate adaptation in pennycress, we performed a climate genome-wide association study (GWAS) on our collection using 19 bioclimatic (BIOCLIM) variables associated with each accession’s collection site. Notably, Eurasian samples exhibited significantly higher h^2^ compared to the North American samples (**Figure 3C**, h^2^_Eurasian_ = 0.853, h^2^_N.American_ = 0.360, *P* = 2.2 × 10^−16^), indicating that climate adaptation in Eurasia is primarily driven by additive genetic variation. In contrast, the reduced h² in North American samples suggests that their dispersal and establishment may have been shaped more by non-genetic factors, such as human-assisted migration, admixture between populations or environmental plasticity.

Using only the Eurasian samples, which exhibited a high h^2^, we performed a climate GWAS on 19 BIOCLIM variables to identify genetic loci underlying climate adaptation. For BIO1 (annual mean temperature), we estimated an h^2^ of 0.94 (s.e. 0.07) and identified 37 significant loci after implementing a Bonferroni correction (**Figure 3D**). Notably, the BIO1 GWAS peaks on chromosomes 2 and 6 overlapped with the peaks from the growth type GWAS, which contained the pennycress gene homologs to *DOG1* and *FLC* in *A. thaliana,* respectively.

Finally, we extended our climate GWAS analysis to all 19 BIOCLIM variables (**Figure S7**) to comprehensively map and document genetic variation associated with diverse climatic conditions for use as a resource for the broader pennycress community.

### The Great Grow-out: field evaluation of our pennycress collection

To assess genetic diversity and agronomic performance within our pennycress collection, we conducted a three-year field experiment, from 2022-2024, evaluating approximately 500 natural accessions under field conditions in Macomb, Illinois, USA (**Figure 4A**). In this paper, we report data from the first two years of the field experiment. Each year, seeds were sown in 4’ x 10’ m plots in the fall, and RGB drone images were collected weekly from November through harvest in 2022-2023, and February through harvest in 2023-2024. All resulting image data was processed to extract canopy coverage, the percentage of ground covered by vegetation, over time. Growth trajectories for each accession was estimated by modeling the total canopy coverage over time, while growth performance was calculated by computing the growth trajectory’s area under the curve (AUC). To assess growth dynamics, we extracted green pixel fractions from images to estimate the progression of maturation and senescence, computing the AUC of green coverage for each accession. We also computed the number of days with canopy coverage above 50% and maximum green coverage for every plot.

**Figure 4.**
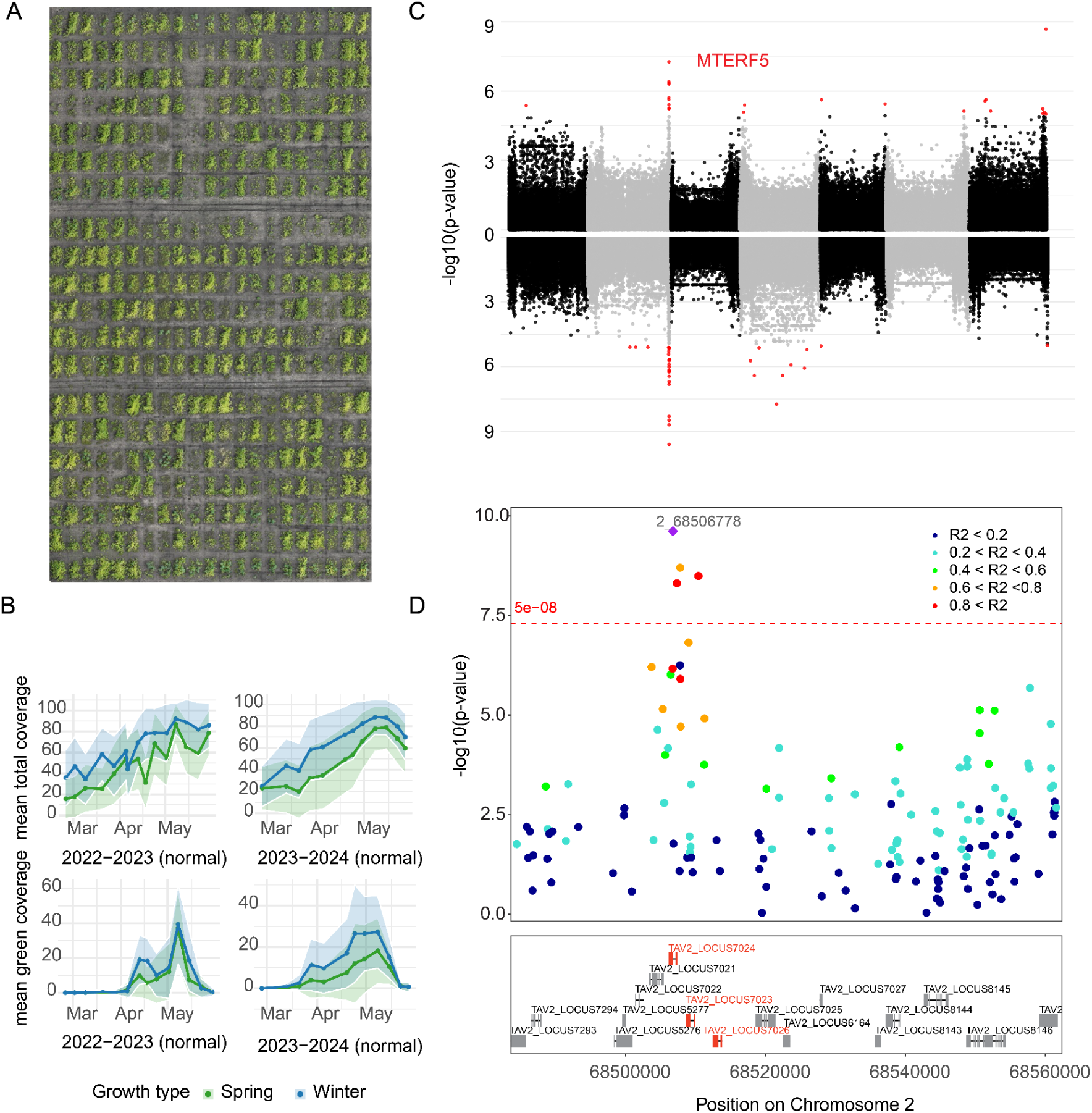
The Great Grow-Out **(A)** The example drone image of part of the great-grow-out field **(B)** The trajectories of total canopy coverage (upper) and green canopy coverage (lower) in two field seasons **(C)** The Manhattan plot of AUC of green coverage trajectories for pennycress accession in the 2022-2023 (upper) and 2023-2024 (lower) seasons. **(D)** LocusZoom plot of the significant peak on chromosome 2; red-highlighted genes represent a cluster of duplicated *mTERF* genes.

Through high-resolution drone image analyses, we found that winter-type pennycress maintained significantly higher canopy coverage compared to spring-type pennycress during both the 2022-2023 and 2023-2024 growing seasons (two-sample t-tests; 2022-2023: *P* = 2.2 x 10^−16^, 2023-2024: *P* = 6.747 x 10^−11^; **Figure 4B, S8**). Overall canopy coverage trajectories were broadly consistent between seasons but exhibited minor differences during the later growth stages. In both years, canopy coverage increased steadily throughout spring, peaking around mid-May. Strikingly, during the 2023-2024 season, we observed a pronounced decline in canopy coverage toward the end of the growing period. The seasonal variation was particularly evident in green coverage trajectories (**Figure 4B)**. In addition, spring-type accessions exhibited significantly lower maximum green area coverage compared to winter-type accessions in 2023-2024, whereas this difference was less pronounced in 2022-2023 (two-sample t-tests; 2022-2023: *P* = 0.020, 2023-2024: *P* = 4.99 × 10^−7^; **Figure S9**). Nevertheless, our data shows winter-type pennycress outperforms the spring-type in the United States Midwest.

### The genetic basis of canopy coverage

To elucidate the genetic determinants underlying growth trajectories in our extensive field experiment, we conducted GWAS for several phenotypes measured during the 2022–2023 and 2023–2024 seasons. These traits included the area under the curve (AUC) for total and green canopy coverage, the number of days with canopy coverage exceeding 50%, and maximum green cover. For the AUC of green canopy coverage, h^2^ was estimated at 0.42 (+/− 0.09) in 2022–2023 and 0.49 (+/− 0.08) in 2023–2024. Notably, both GWAS analyses consistently identified a significant region on chromosome 2, with the lead SNP positioned at 68,507,354 bp. Following LD clumping (r^2^ < 0.5), the associated interval spans 112 kb and encompasses 15 genes. Intriguingly, the lead SNP is located within TAV2_LOCUS7024, in the first intron, a gene that is part of a three-gene tandem duplication cluster; all three genes are homologous to the mitochondrial transcription termination factor (*mTERF*) family in *A. thaliana*. The alternative allele was associated with increased green canopy coverage, implying a prolonged green period during the growing season. Moreover, this same peak was significantly associated with maximum green cover in 2022–2023, as well as with AUC of total canopy coverage, days with canopy coverage above 50%, and maximum green cover in 2023–2024. These findings highlight the role of *mTERF* during vegetative growth and its potential for utilization in future breeding efforts.

## Discussion

Collectively, our comprehensive genomic resource for pennycress greatly enhanced our ability to investigate population dynamics, adaptation history, and the genetic basis underlying agronomically and evolutionarily important traits in pennycress. Leveraging the statistical power gained from studying the largest population of pennycress to date, we first explored pennycress’ complex evolutionary history. The genetic structure and genetic differentiation analyses revealed significant divergence among geographically isolated groups. Notably, we observed a clear separation between samples collected in Asia from those in Europe, and high genetic differentiation of the Armenian (red) and Chinese (purple) subpopulations. These subpopulations are generally geographically separated, as noted previously ^23,27,28,40^, and likely have been evolving in isolation. Intriguingly, we found a subset of North American samples that cluster tightly within the Armenia and Chinese groups. The low genetic differentiation between these North American samples and their Asian counterparts suggests very recent migration events to North America.

The European and North American collected samples represent a much more complex evolutionary history; resulting in 2 separate ancestries. North American samples from Clade 1 likely recently migrated to North America given they are not genetically differentiated from their Eurasian counterparts and that they have the lowest N_e_ out of all sub-clades in the recent past. SFS-based methods to estimate N_e_ are only reliable in the very recent past, <30 generations^41^, nevertheless, North American Clade 1 still has the lowest N_e_. Conversely, the North American samples from Clade 2 likely have had more time to diversify since their initial migration, given they are more genetically differentiated from their Eurasian counterparts and that we were able to identify a candidate selective sweep region containing a gene cluster of MAFs, previously been shown to regulate flowering time in a similar fashion to *FLC*^35,36^. In addition to the multiple inferred migrations to North America, some samples exhibited significant admixture. The presence of individuals in North America with both green and orange subpopulation identities, the identification of a hybrid zone in the Midwestern US, and the low genetic differentiation generally among North American samples collectively points to multiple admixture events. Furthermore, evidence of admixture was also detected among our European samples—specifically between the blue and yellow sub-groups—indicating that admixture is common, despite pennycress having a low genome-wide heterozygosity and a preference to self-fertilize.

Once we had investigated the population history of our pennycress population, we leveraged its standing genetic diversity to map the genetic architecture of key agronomic traits, such as growth type and preferred climatic conditions. In the growth type GWAS, we identified two significant peaks encompassing the *A. thaliana* homologs of *DOG1* and *FLC*. The identification of these homologs in pennycress suggests a conserved genetic framework underlying growth type divergence and flowering time regulation. Notably, a previous study identified four loss-of-function allelic variants of *FLC* associated with the spring growth habit in natural populations, indicating that this trait has arisen multiple times independently^42^. Put together, our improved understanding of the genetic architecture of growth type in pennycress is crucial for breeding, as different climatic regions will require different life cycle strategies.

In our climate GWAS using 19 BIOCLIM variables, we observed a substantial decrease in heritability of the North American samples. This finding aligns with our population history analyses, suggesting that North American pennycress has not yet undergone significant adaptation to the local climate following its migration from Eurasia. Since pennycress is primarily a winter annual with a long day flowering habit that reproduces in the spring, and if this type of climate is very common and similar across temperate regions of the world, pennycress may not need to undergo extensive regional adaptation to succeed in nature. In contrast, the high heritability of the Eurasian samples provides an opportunity to dissect the genetic architecture of climate adaptation in pennycress. We identified several strong association peaks with annual mean temperature, including regions harboring the flowering time regulators *DOG1* and *FLC*, reinforcing the important role of flowering mechanisms in climate adaptation. *FLC* is MADS-box transcription factor that represses flowering in *A. thaliana*^43^, while *SDC* is a F-box factor that plays a key role in epigenetic regulation and circadian clock fine-tuning^44,45^. Since we were able to detect *DOG1* and *FLC* in both the growth type GWAS and the climate GWAS, this further confirms that growth type is a key determinant of climate adaptation, likely through its role in flowering time regulation and seasonal phenology shifts.

Our multi-year field evaluation of natural pennycress accessions provided critical insights into the genetic basis of growth trajectories under field conditions. High-throughput phenotyping using drone imagery revealed substantial variation in canopy coverage and green area dynamics, distinguishing winter- and spring-type accessions. The consistency of growth patterns across seasons underscores the robustness of our phenotyping approach, while seasonal variations highlight the importance of multi-year trials in assessing environmental effects on agronomic performance. Notably, winter-type accessions consistently maintained higher canopy coverage, demonstrating their potential agronomic advantages in temperate climates and promise for use as a winter-time cover crop in between crop rotations. Above all, this comprehensive high-throughput phenotyping approach provides an essential genomic resource to advance pennycress improvement and enables the integration of remote sensing with breeding pipelines.

Finally, our field-based GWAS analysis identified a major locus on chromosome 2 associated with prolonged green coverage. One homolog of the *A. thaliana mTERF* family has been previously implicated in abiotic stress responses, including cold, salt, and metal ion tolerance^46^, which may play a similar role in pennycress. The alternative allele, linked to extended canopy coverage, represents a promising target for breeding climate-resilient pennycress varieties.

In conclusion, this study provides key insights into the evolutionary history of natural pennycress populations and the genetic architecture of important ecological and agricultural traits. We recognize that pennycress’s ecological role differs from its use as an agricultural crop, and the genetic variation influencing traits like canopy coverage may not directly relate to the variation that enhances its competitive ability in natural ecosystems. Although short read sequencing was utilized for this project, future studies should focus on long-read sequencing technologies and pangenome approaches to fully quantify structural variation and minimize reference biases. Nevertheless, these findings pave the way for developing climate-resilient pennycress varieties and the broader adoption of pennycress in agricultural systems.

## Material and Methods

### Plant material collection and climate information

The diversity panel in this study comprised 739 natural accessions of *T. arvense*, including 361 from North America, 318 from Europe, 52 from Asia, and one from South America, with geographic coordinates recorded for each accession. Of these, 479 accessions were collected in this study, while 260 were obtained from previously published datasets, including 13 accessions from Nunn *et al.,* 40 Chinese accessions from Geng *et al.,* and 207 European accessions from Dario *et al.*^28–30^. To characterize the climatic conditions of each accession’s collection site, we retrieved 19 temperature and precipitation BIOCLIM climatic variables^47^ from the WorldClim 2.1 database with 0.5m/s resolution^48^ based on each accession’s geographic coordinates.

### Sequencing and discovery of genomic variations

The resequencing data for 260 previously published accessions were retrieved from the National Center for Biotechnology Information (NCBI) database. We followed a benchmarked plant variant calling pipeline for 739 natural pennycress accessions^31,49^. Low-quality bases and sequencing adapters were trimmed from paired-end reads using Trimommatic v0.33^50^ with parameters ILLUMINACLIP: TruSeq3-PE-2.fa:2:30:10:2:True LEADING:3 TRAILING:3 MINLEN:36. The quality of the trimmed reads was evaluated by FastQC v0.12.1 (https://www.bioinformatics.babraham.ac.uk/projects/fastqc/). Filtered reads were aligned to the pennycress reference genome MN106^29^ using the Burrows-Wheeler Aligner^51^. Alignment sorting and marking duplicated reads were performed using SAMtools^52^ and Picard v3.1.1 (https://broadinstitute.github.io/picard), respectively.

Variant discovery was conducted using the HaplotypeCaller and GenotypeGVCFs functions in GATK with default parameters^53–55^. To remove false positive variants, hard filtering was applied to raw SNP set using GATK with the following parameters ‘QD < 5.0 || FS > 20.0 || MQ < 45.0 || SOR > 2.0 || MQRankSum < −3.0 || ReadPosRankSum < −2.0 || ReadPosRankSum > 2.0’. Additionally, BCFtools was used to remove SNPs with excess heterozygosity^56^. Hard filtering for raw indels was performed using GATK with the following parameters ‘FS > 200.0 || QUAL < 30 || ReadPosRankSum < −20.0 || ReadPosRankSum > 20.0’. Missing genotype imputation and phasing were performed on bi-allelic SNPs with a missing genotype rate below 10% using Beagle v5.3^57,58^. Following imputation, SNPs with a minor allele count below 10 were removed to generate a high-confidence SNP dataset for pennycress.

### Population structure and LD decay analyses

To minimize bias caused by linkage disequilibrium (LD) when estimating the population structure of 739 pennycress natural accessions, we first performed LD pruning to identify independent SNPs for analysis. This was done using PLINK v1.9 with the parameter “-indep-pairwise 2000 100 0.1”, resulting in a total of 11,115 SNPs. Principal component analysis (PCA) was then conducted using the “prcomp” function in R version 4.4.0 based on the LD-pruned SNP set. Identity-by-state (IBS) and hierarchical clustering analyses were performed on LD-pruned SNPs using the R package SNPrelate^59^. To identify subpopulations and assess admixture patterns, we ran ADMIXTURE v1.3.0^60^ on LD-pruned SNPs with values of K ranging from 2 to 20, selecting the optimal K based on cross-validation error.

We estimated the rate of linkage disequilibrium (LD) decay across 739 pennycress accessions, within each sub-clade, and within each subpopulation using PopLDdecay^61^ with the parameter “-MaxDist 500 −MAF 0.01”. To calculate LD decay within subpopulations, we selected individuals primarily assigned to a single subpopulation, defined as having >60% ancestry proportion according to the ADMIXTURE output.

### Growth type, climate, and field experiment GWAS

We conducted GWAS on three traits: growth type, climate variables, and canopy coverage in the *Great Grow-Out* field experiment. For growth type, we evaluated 419 accessions under field conditions, categorizing them as spring or winter based on inflorescence development. The genotype dataset was filtered to remove SNPs absent in these accessions, with a minor allele frequency (MAF) threshold of 0.02. Linkage disequilibrium (LD) pruning (r² < 0.9) was applied using PLINK with parameters −-indep-pairwise 2000 100 0.9, resulting in 414,115 SNPs. For climate GWAS, we analyzed 19 BIOCLIM variables separately for Eurasian and North American samples, applying the same MAF filter and LD pruning, yielding 333,588 SNPs. For the *Great Grow-Out* field experiment GWAS, we examined four drone-derived canopy traits: (1) AUC of total canopy coverage, (2) AUC of green canopy coverage, (3) days with over 50% coverage, and (4) maximum green canopy coverage. After applying the same filtering criteria, 330,668 SNPs remained. We conducted GWAS using two approaches: GEMMA, a SNP-based linear mixed model, and HapFM, a haplotype-based causal inference model. Both methods were used across all analyses.

### Population genetic analyses

The fixation index, or F_ST_, was calculated to assess the level of genetic differentiation among the 2 clades and 7 sub-populations^62^. After LD pruning, yielding roughly 11,000 SNPs across the genome, we used the VCFtools −-weir-fst command to calculate F_ST_ values for every pairwise combination of the four sub-clades and seven sub-populations. To establish 95% confidence intervals for F_ST_ estimates, we performed 1,000 bootstrapping iterations, selecting 20 individuals from each population and recomputing F_ST_ on these subsets.

Tajima’s D was calculated across 5.4 million SNPs genome-wide to investigate the demographic history of pennycress^32^. The VCFtools −-TajimaD command was used with a bin size of 10 kb.

XP-CLR was utilized to detect selective sweeps within the two main pennycress clades^34^. A weighted CLR was used to downweight SNPs in high LD (r^2^ > 0.95), a phased genotype matrix along with a genetic map was used as input with a spacing of 2kb, sliding window of 0.5 cM, and a minimum 200 SNPs per window.

LDhat was used to generate a genetic map of the entire pennycress collection^63^. 96 diploid pennycress accessions were randomly sampled and used for genetic map creation. We used the pre-computed look-up table for Θ = 0.001 and N = 192 for genetic map estimation.

The proportion of heterozygous sites is used to uncover the degree of inbreeding within a population. PLINK −-het was utilized to count the number of heterozygous sites of individuals across the 4 sub-groups and the 7 sub-populations of pennycress^64^.

Stairway Plot 2 was used to infer recent demographic history of pennycress sub-clades and subpopulations^33^. Since Stairway Plot 2 requires a SFS as input, we inferred a folded SFS using vcf2sfs, which converts VCF files into the SFS in the fastsimcoal2 format^65^. For each subpopulation, 200 random subsamples from each sub-group were used to train the model. Four breakpoints were used at 1/5, 2/5, 3/5, and 4/5 positions of the maximum possible breakpoint, based on the number of sequences in each subpopulation. These breakpoints define potential changes in population size throughout the demographic history. Additionally, we used a proxy mutation rate of 7e^-9^ mutations per site per generation based on the mutation rate for *A. thaliana*^66^, assuming a generation time of 1 year. We report the median of the N_e_ estimates (**Figure 2**) as well as create pseudo confidence intervals from the upper (97.5%) and lower (2.5%) estimates.

### Field experiment

Field trials were conducted over two growing seasons (2022–2023 and 2023–2024) at Western Illinois University in Macomb, IL, USA, at coordinates 40.4910, −90.6870 and 40.4920, −90.6885, respectively. Different fields were used for the two years due to avoid issues of seed shattering and persistence from previously planted wild black seeded pennycress plots. For the 2022-2023 field trial, the preceding crop was soybean followed by oats for silage harvested in mid June. For the 2023-2024 field trial, the preceding crop was soybean followed by annual rye harvested for silage in mid June. In both trials, seeds were drilled into 7 rows spaced 15cm apart with plot sizes of 4’ x 10’. Planting dates for fall 2022 were October 1-2, and September 30-Oct 1 for fall 2023. Across both years, plots were maintained by hand weeding and grass herbicides as needed.

### Drone image data processing

For 2022-23, the first drone images were taken on 11/02/2022, and the final images for the season were taken on 05/29/2023. The entire field was imaged on average every 7 days, with a total of 30 flights. Excluding flight days that were covered in snow, the maximum gap between imaging dates was 18 days. For fall 2022, images were taken with a DJI AIR 2S with 20 MP camera at a height of 20m. In the spring of 2023, we moved to DJI MAVIC 3 with a multispectral 20 MP camera. For the 2023-2024 growing season, the first drone images were taken on 02/27/2024, and the final images for the season were taken on 05/27/2024. The entire field was imaged on average every 8 days, with a total of 12 flights. Excluding flight days that were covered in snow the maximum gap between imaging dates was 15 days. Flights were pre-programmed using DJI Pilot 2 software and conducted at a height of 17 meters above ground level, resulting in an image resolution of approximately 0.75 cm/pixel. The drone maintained a flight speed of ∼1.5 m/s, with 70% side overlap and 80% front overlap between images to ensure sufficient coverage for stitching. All flights were conducted around 9:00 AM CDT to maintain consistency in lighting. Captured images were processed into orthomosaics using WebODM (version 2.5).

Orthomosaic images were analyzed using a Python script developed with OpenCV libraries^67^. A reference scale was established for each orthomosaic using known field dimensions, and a scaling factor was applied to ensure consistent spatial resolution across plots. Orthomosaic images were then rotated to standardize orientation prior to analysis. Individual plots were automatically cropped from the orthomosaics based on a predefined grid layout. To distinguish vegetation from background soil, a soil mask was generated using the Blue-Green Index (BGI), where pixels with BGI < 0.06 were classified as soil. Additionally, vegetation status was assessed using the Green Leaf Index (GLI) to separate green from senesced plant tissue. A GLI threshold of 0.2 was used to classify vegetation with GLI ≥ 0.2 classified as green, and GLI < 0.2 classified as senesced. The proportions of green and senesced vegetation were calculated as a percentage of total non-soil pixels within each plot. A set of reference images was generated for each plot by overlaying the BGI and GLI masks onto the original image for quality control of plot boundaries and index thresholds. The vegetation indices were calculated as: BGI=(G-B)/(G+B+ε), and GLI=(2*G-R-B)/(2*G+R+B+ε), where R, G, and B represent the red, green, and blue channels of each image, respectively, and ε is a small constant added to avoid division by zero.

## Data availability

All raw sequencing data generated in this study have been deposited in the NCBI Sequence Read Archive under BioProject accession number PRJNA1239920. Publicly available whole-genome resequencing datasets from previous studies were obtained from NCBI BioProject PRJNA715950 and the ENA Sequence Read Archive under study accession numbers PRJEB50950 and PRJEB46635.

## Acknowledgements

We thank the numerous citizen scientists who graciously collected *Thlaspi arvense* seeds and posted their sightings on iNaturalist. We thank iNaturalist for creating a user-friendly and impactful platform for casual observers, citizen scientists, and professional scientists alike. Special thanks to Patrick Bulman, Markéta Machová, Julia Proulx, and Dan Wrench for serving as hubs and forwarding/donating seeds with the necessary permits. We thank the Arabidopsis Biological Resource Center (ABRC) and the Nottingham Arabidopsis Stock Center (NASC) for curating and distributing related seeds.

## Supplementary Tables

**Table S1.** Meta information about the 739 pennycress natural accessions

Meta information, including the geographic coordinates, country, growth type and data source of 739 pennycress accessions used in this study.

Table_S1_accession_meta_information.txt

**Table S2**. Summary of SNPs and Indels identified from 739 pennycress natural accessions

The summary of count and estimated functional importance of SNPs and Indels identified from 739 pennycress natural accessions

Table_S2_SNP_Indel_summary.txt

## Supplementary Figures

**Figure S1.**
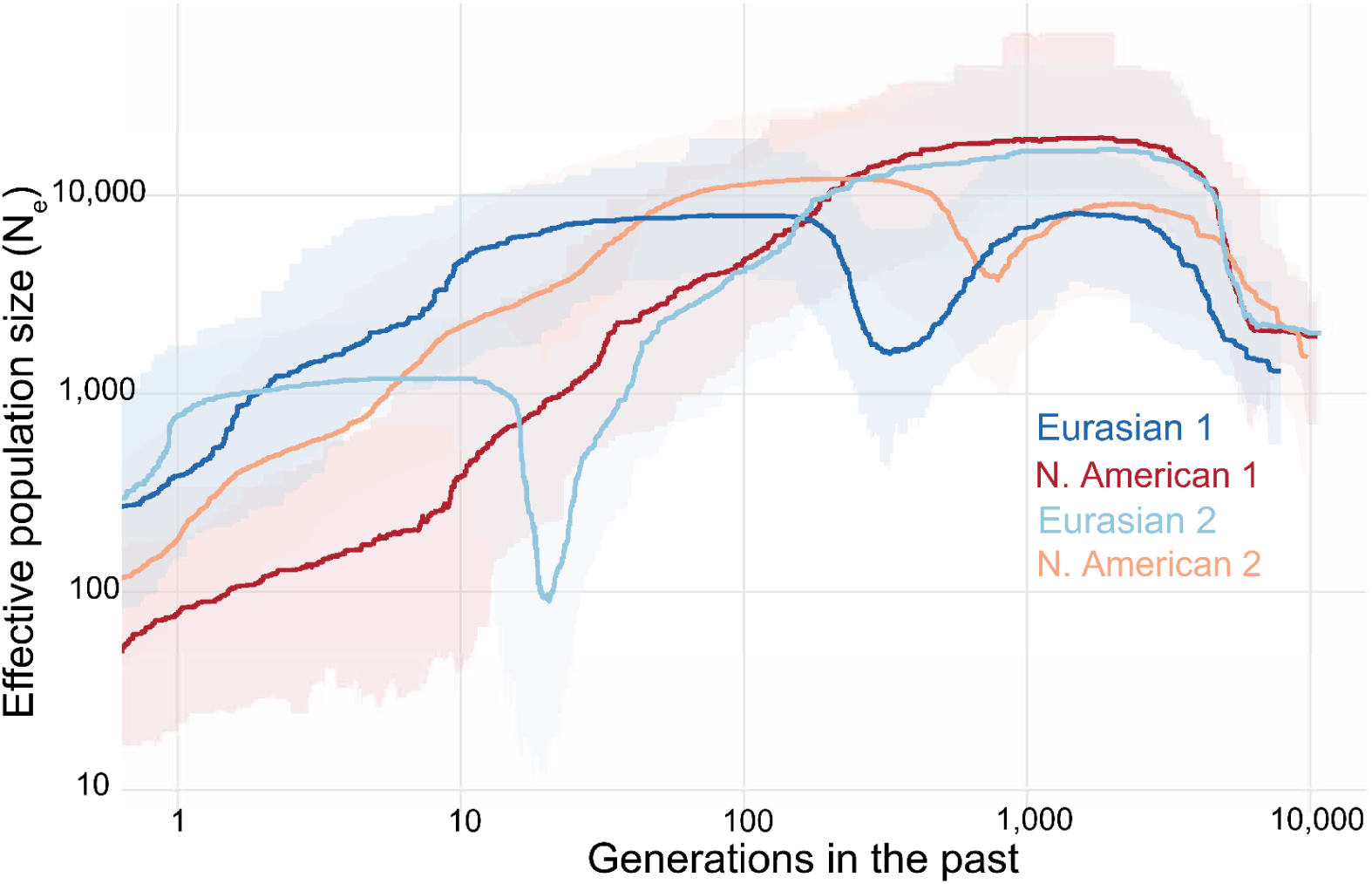
Effective population size estimates in the recent past for Eurasian and North American samples from Clades 1 and 2 based on the SFS.

**Figure S2.**
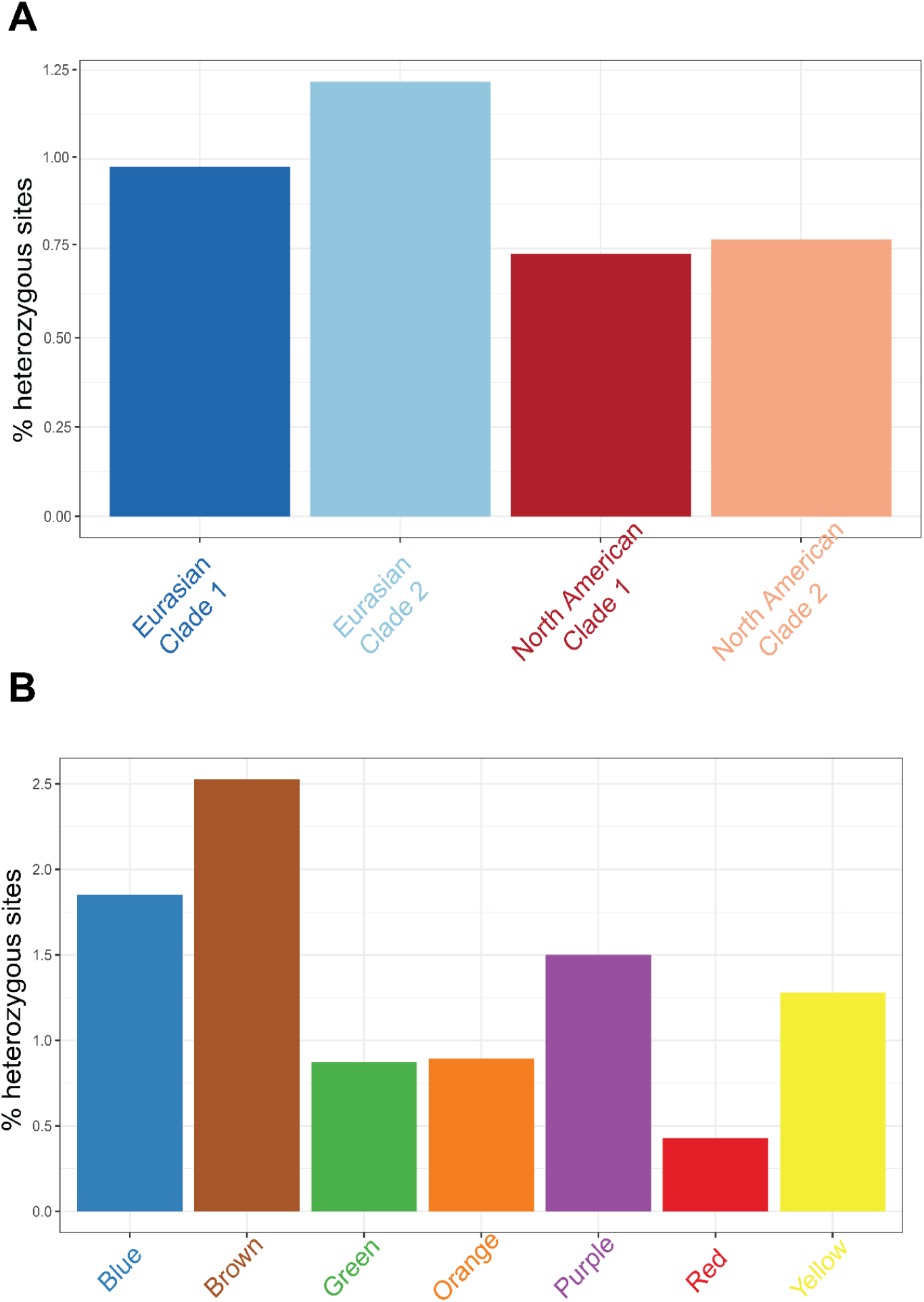
Barcharts of the percentage of heterozygous sites in pennycress. **(A)** The heterozygosity of Eurasian and North American samples in Clades 1 and 2 based on IBS genetic distance and **(B)** the heterozygosity of the STRUCTURE-defined seven subpopulations.

**Figure S3.**
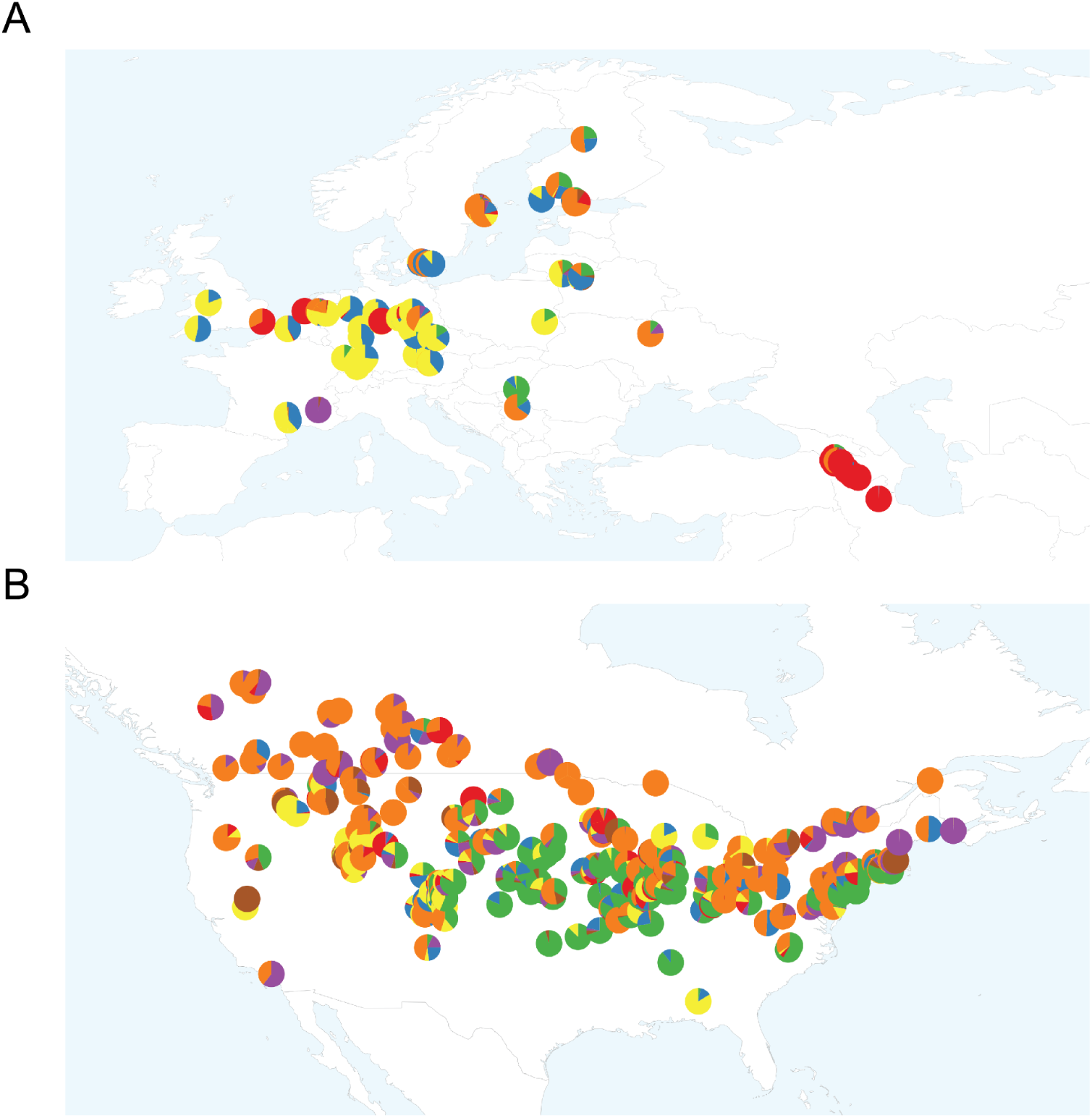
Zoom-in map of accession collection sites with STRUCTURE subpopulation inference in **(A)** Europe and **(B)** North America. The color corresponds to the proportions of each subpopulation estimated by STRUCTURE (*K*=7).

**Figure S4.**
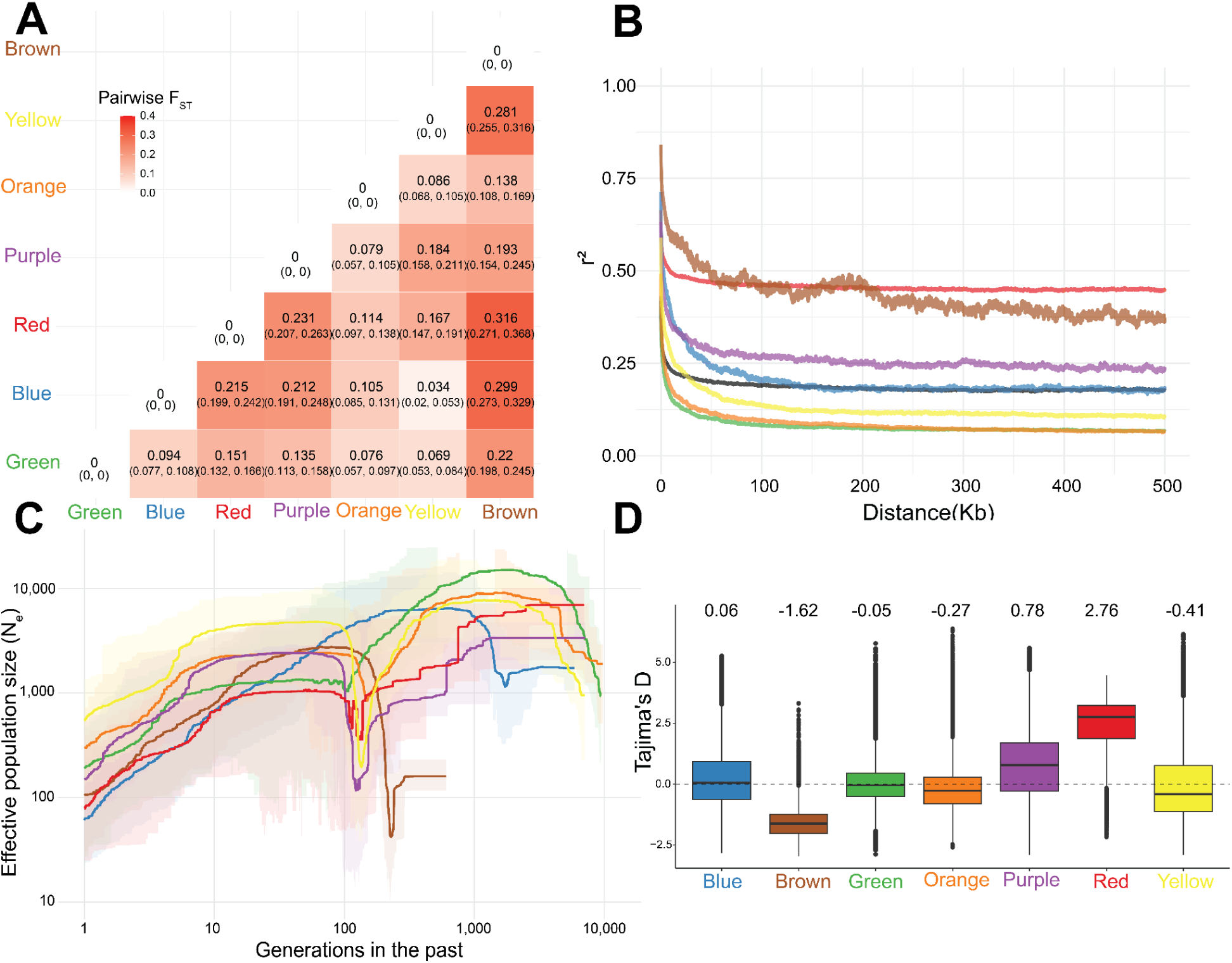
The history and dynamics of seven subpopulations identified by STRUCTURE. **(A)** Pairwise F_ST_ between the seven subpopulations identified by STRUCTURE. The values are median genome-wide F_ST_ between two populations with a 95% confidence interval. **(B) T**he rate of LD decay in seven subpopulations. The black line indicates the average rate of LD decay in all 739 pennycress natural accessions. **(C)** Effective population size estimates in the recent past for the seven subpopulations. **(D)** Boxplots of Tajima’s D calculated in 10kb windows genome-wide for each subpopulation.

**Figure S5.**
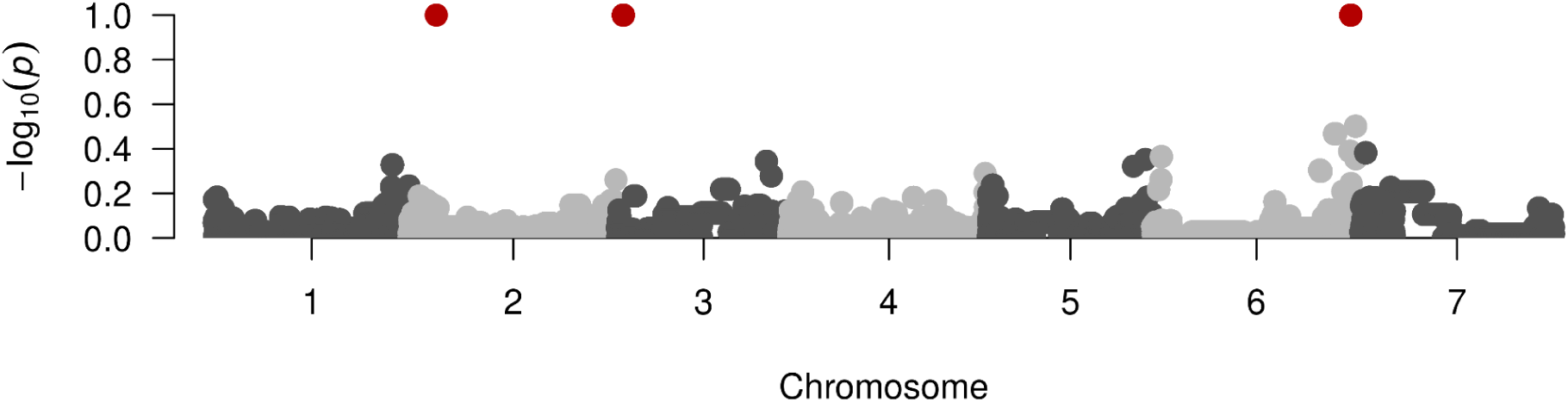
Manhattan plot of HapFM results on growth-type. The y-axis represents the posterior inclusion probability of the haplotype block. The red dot on chromosome 2 represents *DOG1,* red dot on chromosome 3 marks *SDC*, and the red dot on chromosome 6 is *FLC*.

**Figure S6.**
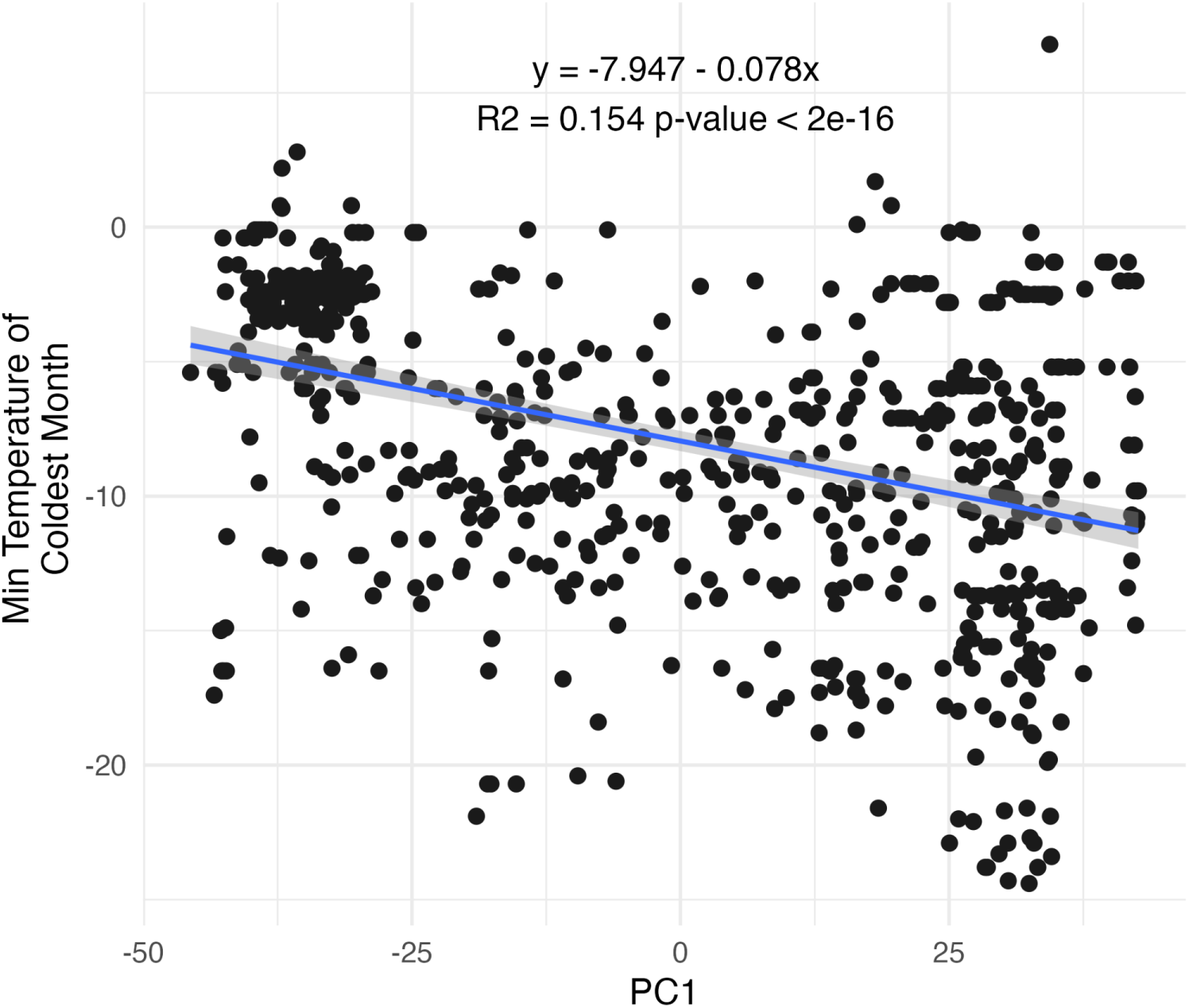
Association between PC1 from the genetic principal component analysis and the minimum temperature of the coldest month at pennycress collection sites.

**Figure S7.**
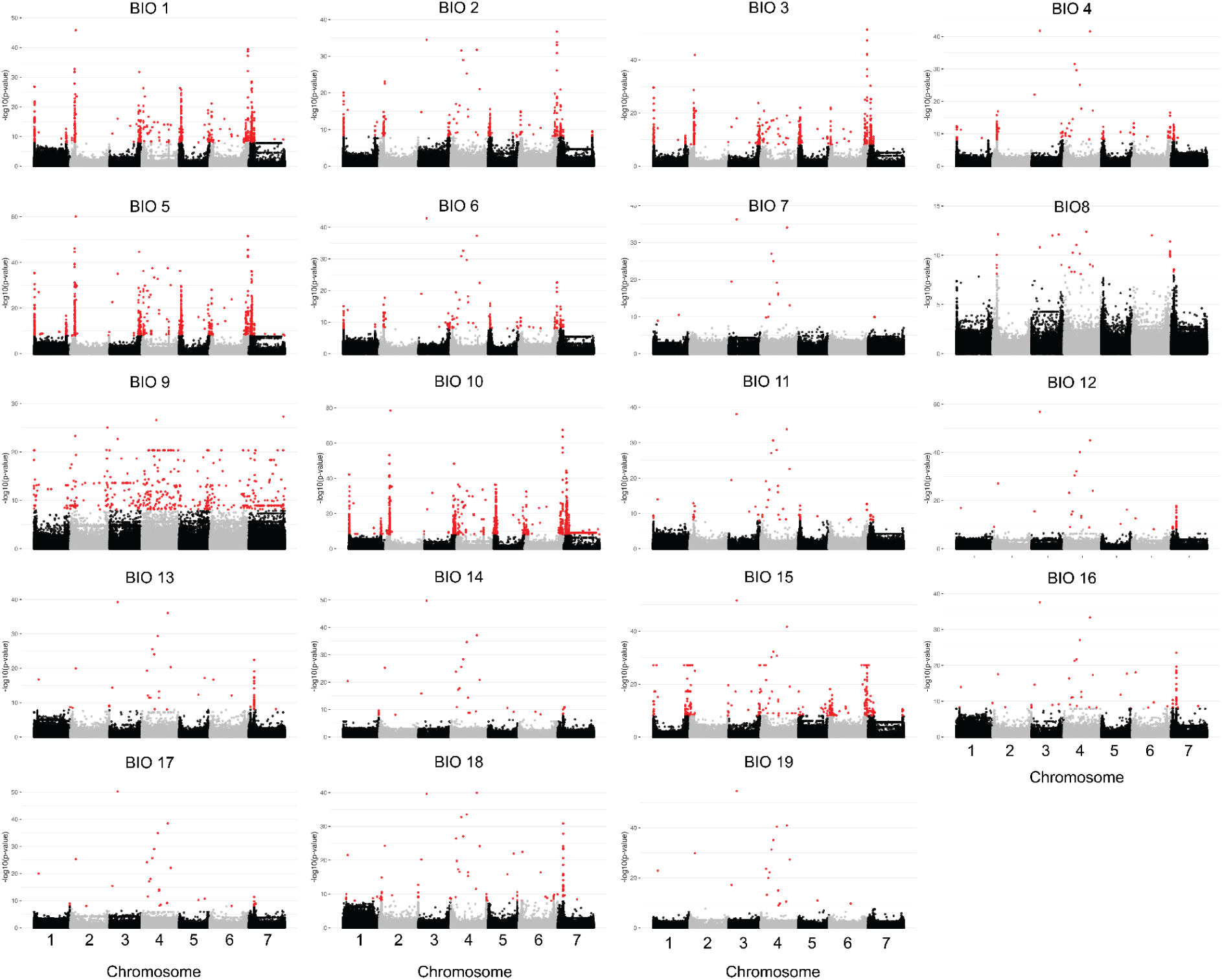
Manhattan plot showing the results from GWAS of the 19 BIOCLIM climate variables in the Eurasian pennycress population. The y-axis represents the −log₁₀ *p*-value for each SNP. Red points indicate SNPs that passed the Bonferroni correction threshold.

**Figure S8.**
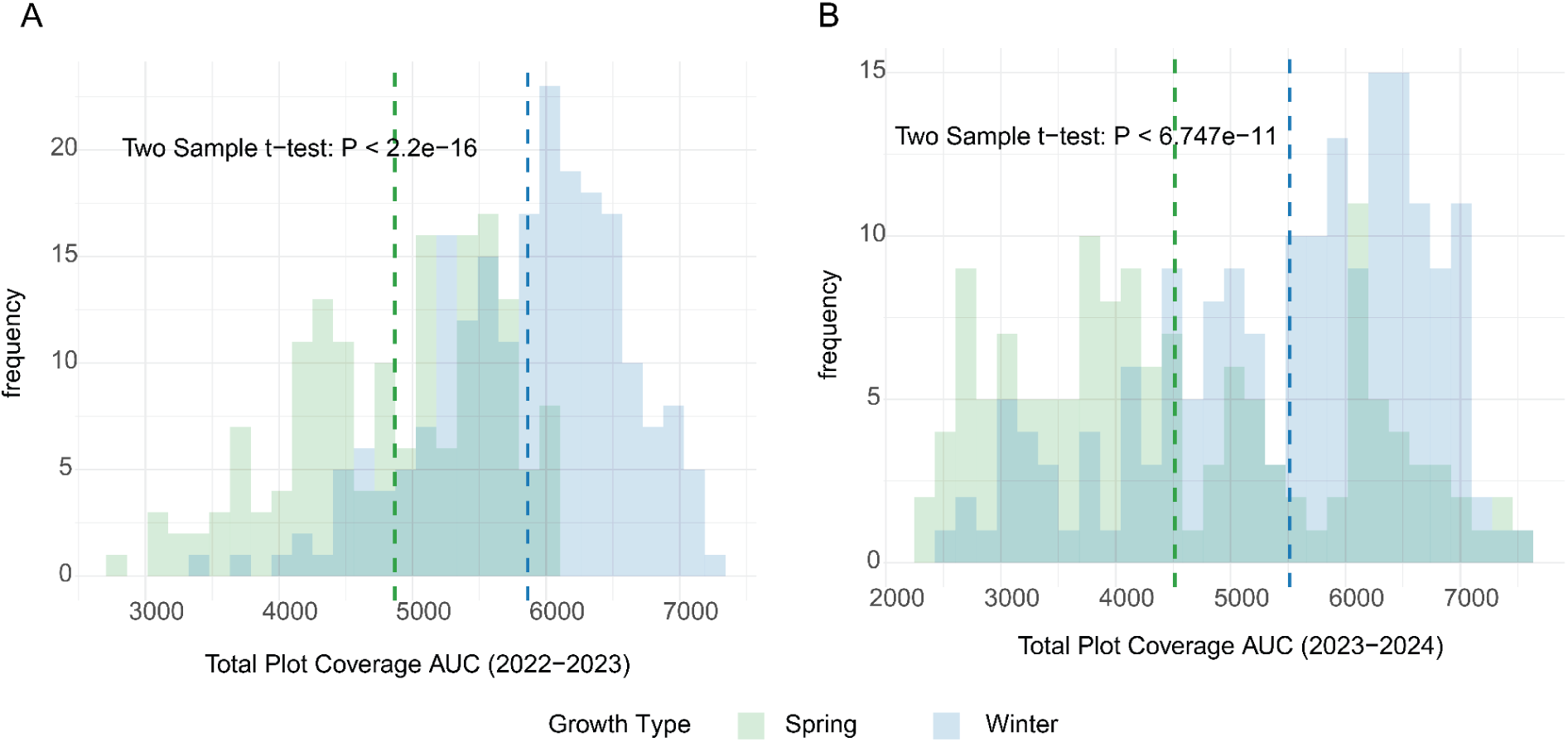
Histogram of the Area Under the Curve (AUC) for total plot coverage of each pennycress accession in the Great Grow Out field experiment during the 2022–2023 and 2023–2024 seasons. Spring and winter growth types are indicated by green and blue colors, respectively.

**Figure S9.**
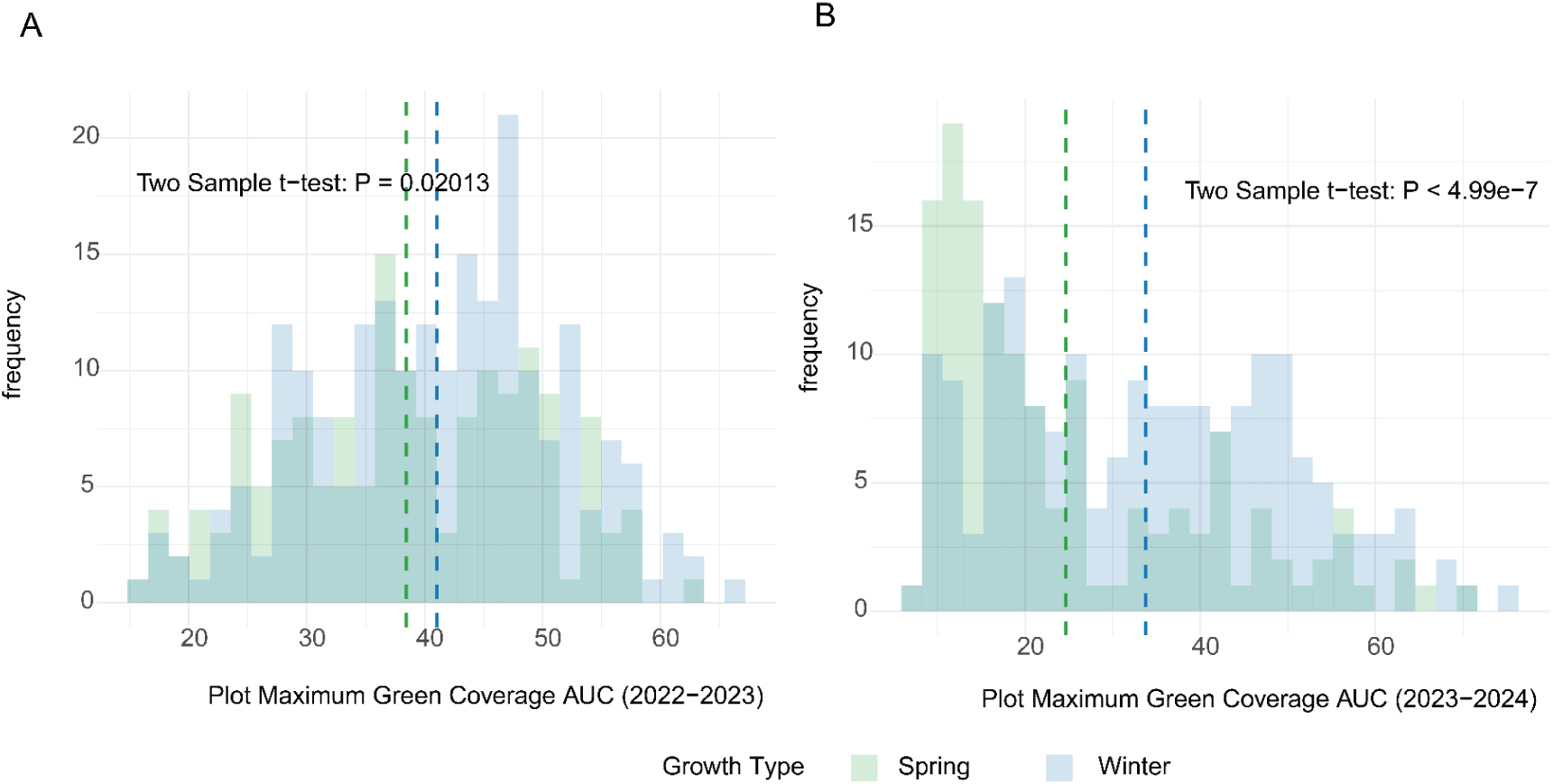
Histogram of the maximum green coverage of each pennycress accession in the Great Grow Out field experiment during the 2022–2023 and 2023–2024 seasons. Spring and winter growth types are indicated by green and blue colors, respectively.

